# Transcription Co-Factor LBH Is Necessary for Maintenance of Stereocilia Bundles and Survival of Cochlear Hair Cells

**DOI:** 10.1101/2020.05.13.093377

**Authors:** Huizhan Liu, Kimberlee P. Giffen, Grati M’Hamed, Seth W. Morrill, Yi Li, Xuezhong Liu, Karoline J. Briegel, David Z. He

## Abstract

Hearing loss affects ~10% of adults worldwide and is irreversible. Most sensorineural hearing loss is caused by progressive loss of mechanosensitive hair cells (HCs) in the cochlea of the inner ear. The molecular mechanisms underlying HC maintenance and loss are largely unknown. Our previous cell-specific transcriptome analysis showed that Limb-Bud-and-Heart (LBH), a transcription co-factor implicated in development, is abundantly expressed in outer hair cells (OHCs). We used *Lbh*-null mice to identify its role. Surprisingly, *Lbh* deletion did not affect differentiation and early development of HCs, as nascent HCs in *Lbh* knockout mice had normal looking stereocilia bundles. Whole-cell recording showed that the stereocilia bundle was mechanosensitive and OHCs exhibited the characteristic electromotility. However, *Lbh*-null mice displayed progressive hearing loss, with stereocilia bundle degeneration and OHC loss as early as postnatal day 12. Cell-specific RNA-seq and bioinformatic analyses identified *Spp1, Six2, Gps2, Ercc6, Snx6* as well as *Plscr1, Rarb, Per2, Gmnn* and *Map3k5* among the top five transcription factors up- or down-regulated in *Lbh*-null OHCs. Furthermore, this analysis showed significant gene enrichment of biological processes related to transcriptional regulation, cell cycle, DNA damage/repair and autophagy. In addition, Wnt and Notch pathway-related genes were found to be dysregulated in *Lbh*-deficient OHCs. We speculate that LBH may promote maintenance of HCs and stereocilia bundles by regulating Notch and Wnt signaling activity. Our study implicates, for the first time, loss of LBH function in progressive hearing loss, and demonstrates a critical requirement of LBH in promoting HC survival.

## Introduction

466 million people worldwide are estimated to be living with hearing loss. Most sensorineural hearing loss is caused by progressive degeneration of hair cells (HCs) in the cochlea of the inner ear. These cells are specialized mechanoreceptors which transduce mechanical forces transmitted by sound to electrical activities (Hudspeth, 2014; Fettiplace, 2017). HCs in adult mammals are terminally differentiated and unable to regenerate once they are lost due to aging or exposure to noise and ototoxic drugs. Although HCs have been well characterized morphologically and biophysically, the key molecules that control their differentiation, homeostasis and aging remain to be identified.

Inner and outer HCs (IHCs and OHCs) are the two types of HCs, with distinct morphology and function in the mammalian cochlea (Dallos, 1992). IHCs are the true sensory receptor cells and transmit information to the brain while the OHCs are a mammalian innovation with a unique capability of changing its length in response to changes in receptor potential (Brownell et al., 1985; Zheng et al., 2000). OHC motility is believed to confer the mammalian cochlea with high sensitivity and exquisite frequency selectivity (Liberman et al., 2002; Dallos et al., 2007). We recently compared cell type-specific transcriptomes of IHC and OHC populations collected from adult mouse cochleae to identify genes commonly and differentially expressed in these cells (Liu et al., 2014; Li et al., 2018). Our analysis showed that *Limb-bud-and-heart* (*Lbh)*, a transcription co-factor implicated in development (Briegel & Joyner 2001; Briegel et al., 2005; Ai et al. 2008; Al-Ali et al., 2010; Lindley et al, 2015), is expressed in both IHCs and OHCs (Liu et al., 2014; Li et al., 2018). *Lbh* is also expressed in vestibular HCs (Scheffer et al., 2015) and upregulated during transdifferentiation from supporting cells to HCs (Ebeid et al., 2017; Yamashita et al., 2018). We, therefore, asked whether LBH is necessary for HC differentiation, development, and maintenance. Because *Lbh* expression in OHCs is 4.7 Log2 fold greater than in IHCs, we also questioned whether LBH plays a role in regulating cell specialization underlying OHC morphology and function.

*Lbh* conditional knockout mice have been generated by LoxP and Cre recombination (Lindley and Briegel, 2013). The role of LBH in HCs was examined by comparing changes in morphology, function and gene expression between HCs from *Lbh-*null and wildtype mice. Results showed that HC differentiation, formation of the mechanotransduction apparatus and OHC specialization were unaffected by loss of LBH. However, stereocilia bundles and HCs, especially OHCs, showed signs of degeneration as early as P12. Moreover, adult *Lbh*-null mice displayed progressive loss of hearing and otoacoustic emissions, suggesting that LBH is critical for maintenance of stereocilia bundles and survival of HCs. Cell-specific transcriptome and bioinformatics analyses showed a significant enrichment of genes associated with transcription, cell cycle, DNA damage/repair, and autophagy in the *Lbh*-null OHCs. Wnt and Notch pathway-related genes, known for their important roles in regulating HC differentiation and regeneration in vertebrate HCs (Raft and Groves, 2015), were found to be dysregulated. We speculate that dysregulated Notch/Wnt activity following LBH ablation may lead to degeneration of stereocilia bundles and OHCs. Our study implicates, for the first time, loss of transcription co-factor LBH function in progressive hearing loss, and demonstrates a critical requirement of LBH in promoting cochlear HC survival.

## Results

### 1. Expression of *Lbh*/LBH in inner ear HCs

*Lbh* gene expression in HCs and supporting cells in the adult murine organ of Corti was examined using our published cell type-specific RNA-seq data sets (Liu et al., 2018). *Lbh* was expressed in all four cell types, IHCs, OHCs, pillar cells and Deiters’ cells, however, *Lbh* transcript levels were highest in OHCs (Fig. 1A, left panel). We also examined expression of *Lbh* during development using RNA-seq data by Scheffer et al (Scheffer et al., 2015). *Lbh* was expressed with comparable levels in both cochlear and vestibular HCs at embryonic day 16 (E16) and upregulated at postnatal day 7 (P7) (Fig. 1A; right panel). In contrast, low level *Lbh* expression in non-sensory supporting cells did not change. We next used LBH-specific antibodies to examine LBH protein expression in inner ears from neonatal and adult C57BL/6 mice. Fig. 1B shows a micrograph obtained from a cryosection of a P3 cochlea. LBH was expressed in both OHCs and IHCs with no obvious expression in supporting cells in the organ of Corti at this neonatal stage (Fig. 1B). LBH positivity was also detected in some cells in the greater epithelial ridge. In P12 cochlea, LBH was still expressed in both IHCs and OHCs, as revealed by confocal microscopy, however, expression was strongest in OHCs (Fig. 1C). Of note, in OHCs LBH was predominately cytoplasmic although weaker expression was also seen in the nuclei of these cells, and in IHCs (Fig. 1C). This expression pattern was LBH-specific, as in the age-matched *Lbh-*null mice, no LBH protein was detected in IHCs and OHCs (Fig. 1D). In adult cochlea, strong LBH expression in OHCs persisted, while LBH expression in IHCs remained weak (Fig. 1E). This pattern of expression is consistent with the predominant expression of *Lbh* mRNA in adult OHCs (Liu et al., 2014, 2018). In contrast, LBH was not expressed in vestibular HCs, as no LBH-specific immunopositivity was detected in utricular HCs of P12 wildtype mice (Figs. 1F,G).

**Figure 1:**
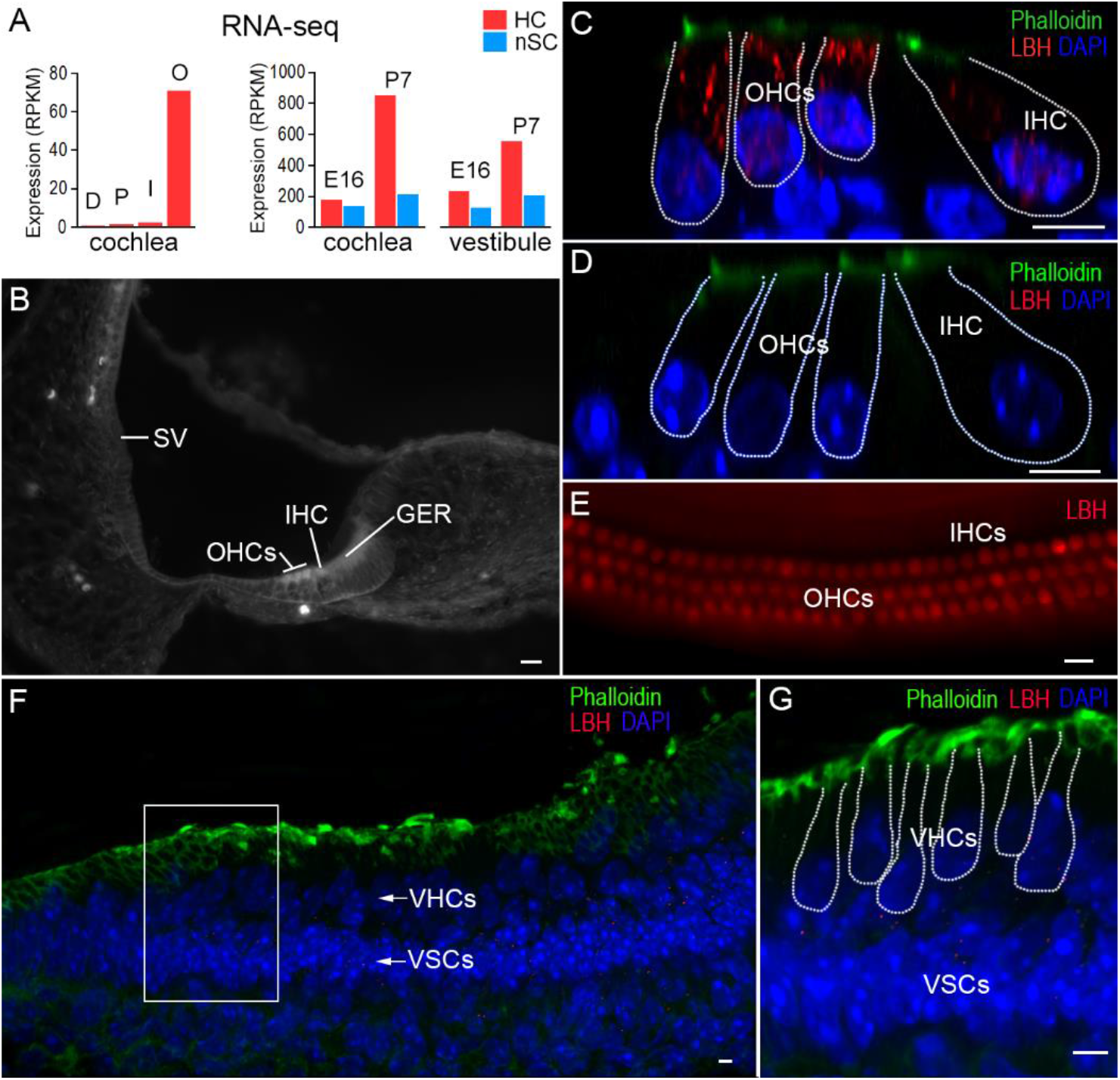
Expression of LBH in cochlear and vestibular hair cells. **A**: Cell type-specific expression of *Lbh* mRNA in Deiters’ cells (D), pillar cells (P), IHCs (I) and OHCs (O), as well as in vestibular hair cells (VHCs), and non-sensory cells (VSCs) during development. **B**: Fluorescent microscopy picture of antibody staining of LBH protein in a cryosection of the cochlea from a P3 wildtype mouse. Stria vascularis (SV), IHCs, OHCs, and greater epithelium ridge (GER) are marked. Bar: 10 μm. **C**: LBH expression (red) in the organ of Corti from a P12 wildtype mouse using optical sectioning with confocal microscopy. **D**: Lack of LBH protein expression in hair cells in a P12 *Lbh*^Δ2/Δ2^ null mouse. Bars: 5 μm in C and D. **E**: Fluorescence microscopy picture of antibody staining of LBH in P30 cochlear hair cells from wildtype mouse. **F** and **G**: Cryosection of the utricle from a P12 wildtype mouse. The nuclei of vestibular hair cells (VHCs) and supporting cells (VSCs) are marked white arrows. The area delineated by the white frame in F is displayed at higher magnification in G. Bars: 5 μm in F and G.

### 2. Auditory function of *Lbh*-mutant mice

To determine if LBH expression in cochlear HCs is required for hearing, we examined auditory function in *Lbh*-mutant mice by measuring auditory brainstem response (ABR). Fig. 2A shows the ABR thresholds of homozygous (*Lbh*^Δ2/Δ2^), heterozygous (*Lbh*^+/Δ2^) and wildtype (*Lbh*^+/+^) mice at 1 month of age. As shown, the threshold of *Lbh*^Δ2/Δ2^ null mice is elevated by ~10 dB at lower frequencies to ~40 dB in higher frequencies relative to their wildtype littermates. Heterozygous *Lbh*^+/Δ2^ mice also showed 10 to 25 dB hearing loss at higher frequencies when compared with the wildtype controls, suggesting that even minor decreases in *Lbh* gene dosage impairs hearing. We next measured distortion product otoacoustic emission (DPOAE) thresholds at 8, 16 and 32 kHz in these mice. DPOAEs are generated by motor activity of OHCs (Liberman et al., 2002; Dallos et al., 2007) and reflect OHC function/condition. Consistent with our ABR measurements, DPOAE thresholds (Fig. 2B) were also elevated at higher frequencies in *Lbh*^Δ2/Δ2^, and *Lbh*^+/Δ2^ mice. We further measured the cochlear microphonic (CM) response to an 8 kHz tone burst in *Lbh*^Δ2/Δ2^ and *Lbh*^+/+^ mice. A significant reduction of the CM magnitude (Fig. 2C) in response to the same level of sound stimulation was observed in *Lbh*^Δ2/Δ2^ mice (n = 6, *p* = 4.29E-06). Since weak expression of *Lbh* was also detected in intermediate cells of the stria vascularis during development (20), we measured endocochlear potential (EP) from one-month-old *Lbh*^Δ2/Δ2^ and *Lbh*^+/+^ mice to determine whether stria development and function are affected by deletion of *Lbh*. This is necessary since stria function (i.e., the EP) can influence HC survival (Liu et al., 2016). An increase in the EP magnitude was observed in *Lbh*^Δ2/Δ2^ mice (Fig. 2D). The fact that no EP reduction was observed suggests that loss of LBH does not affect stria function. Finally, ABR and DPOAE measurements at 3 months of age showed that hearing was further decreased in both *Lbh*^Δ2/Δ2^ and *Lbh*^+/Δ2^ mice (Figs. 2E,F), indicating LBH deficiency causes progressive hearing loss.

**Figure 2:**
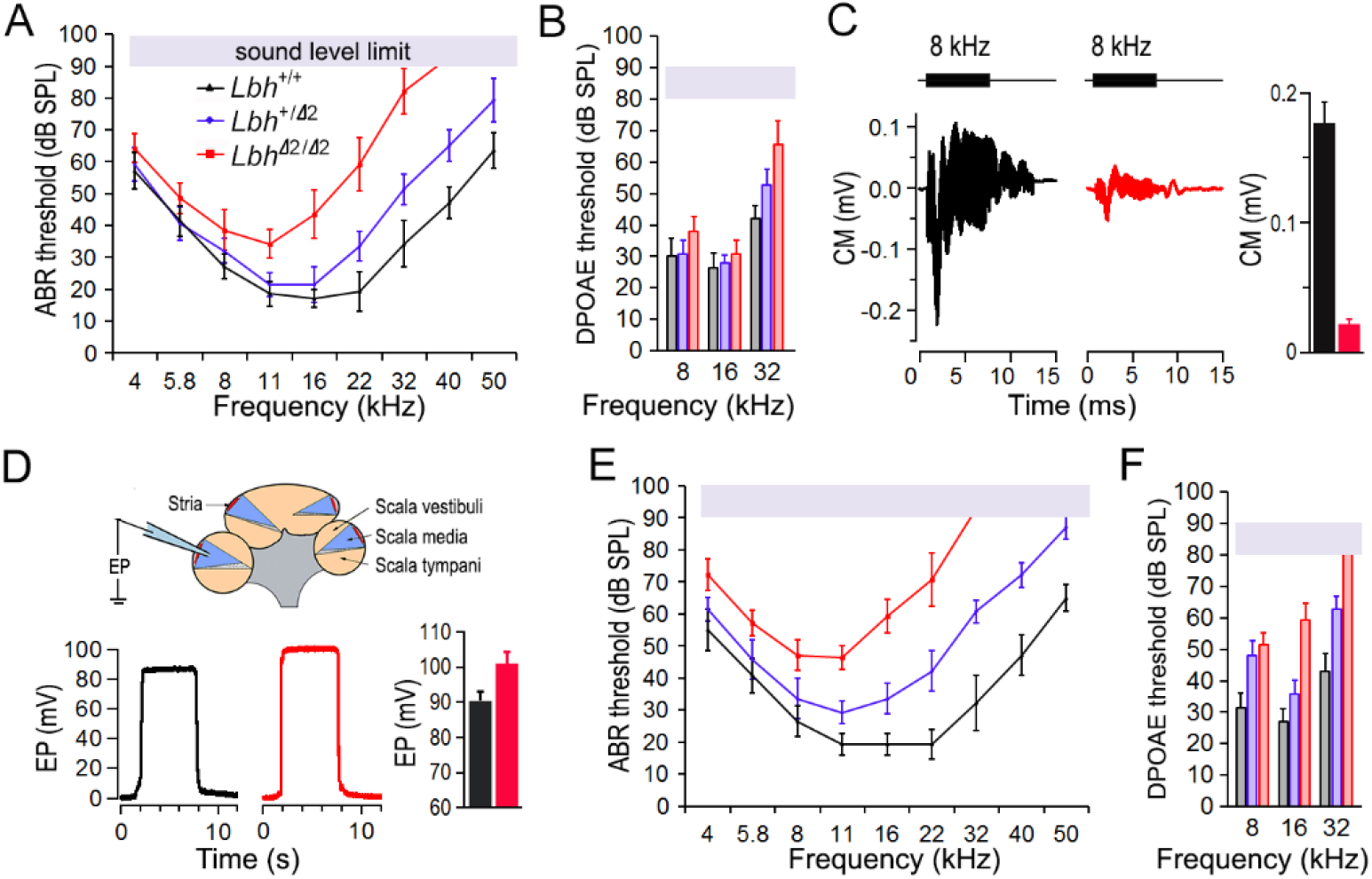
Auditory function of *Lbh*-mutant mice. **A**: ABR thresholds of the three genotypes of mice (color-coded) at 1 month-of-age. Eight mice for each genotype from three different litters were used. **B**: DPOAE thresholds at 1 month. **C**: Representative CM responses together with CAP measured in *Lbh*^Δ2/Δ2^ null (red) and *Lbh*^+/+^ wildtype (black) mice. 8 kHz tone bursts (80 dB SPL) were used to evoke response. Peak-to-peak magnitude (mean ± SD, n=6 per genotype) of the CM is presented in the right panel. **D**: Representative EP measured from *Lbh*^Δ2/Δ2^ (red) and *Lbh*^+/+^ (black) mice at 1 month. EP magnitude (mean ± SD, n = 6 per genotype) is also presented. **E**: ABR thresholds at 3 months. **F**: DPOAE thresholds at 3 months.

### 3. Morphological changes of HCs in *Lbh*^Δ2/Δ2^ mice

We next asked if there was progressive HC loss in *Lbh*-deficient mice. To this end, we examined HC frequency at the base and apex of the cochleae at four different ages in *Lbh*^Δ2/Δ2^ and *Lbh*^+/+^ mice (n=3 each). Fig. 3A shows representative confocal images at P12 and 1 month. The total number of IHCs and OHCs at the two cochlear locations were also counted. Fig. 3B shows the percentage of surviving HCs at P3, P12, 1 and 3 months. No HC loss was apparent at either location in P3 *Lbh*^Δ2/Δ2^ or *Lbh*^+/+^ cochleae. At P12, *Lbh*^Δ2/Δ2^ cochlea exhibited sporadic HC loss in the basal turn region, whereby OHC loss was more severe than IHC loss (Figs. 3A, B). At 1 month, OHC loss also occurred at the apical turn, and nearly 50% of OHCs were lost in the basal turn region of these mice (Figs. 3A, B). IHC loss at the basal turn region remained mild. Finally, more OHCs were lost in both apical and basal turns at 3 months, with only ~10% of OHCs remaining in the basal turn region of *Lbh*^Δ2/Δ2^ KO cochleae (Fig. 3B). Interestingly, the majority of IHCs in the basal turns survived and apical IHCs were unaffected, despite substantial OHCs loss at 3 months. We also examined HC survival in the vestibular end organs at 3 months, but did not find any noticeable HC loss in the utricle and crista ampullaris of *Lbh*^Δ2/Δ2^ KO mice (Fig. 3C).

**Figure 3:**
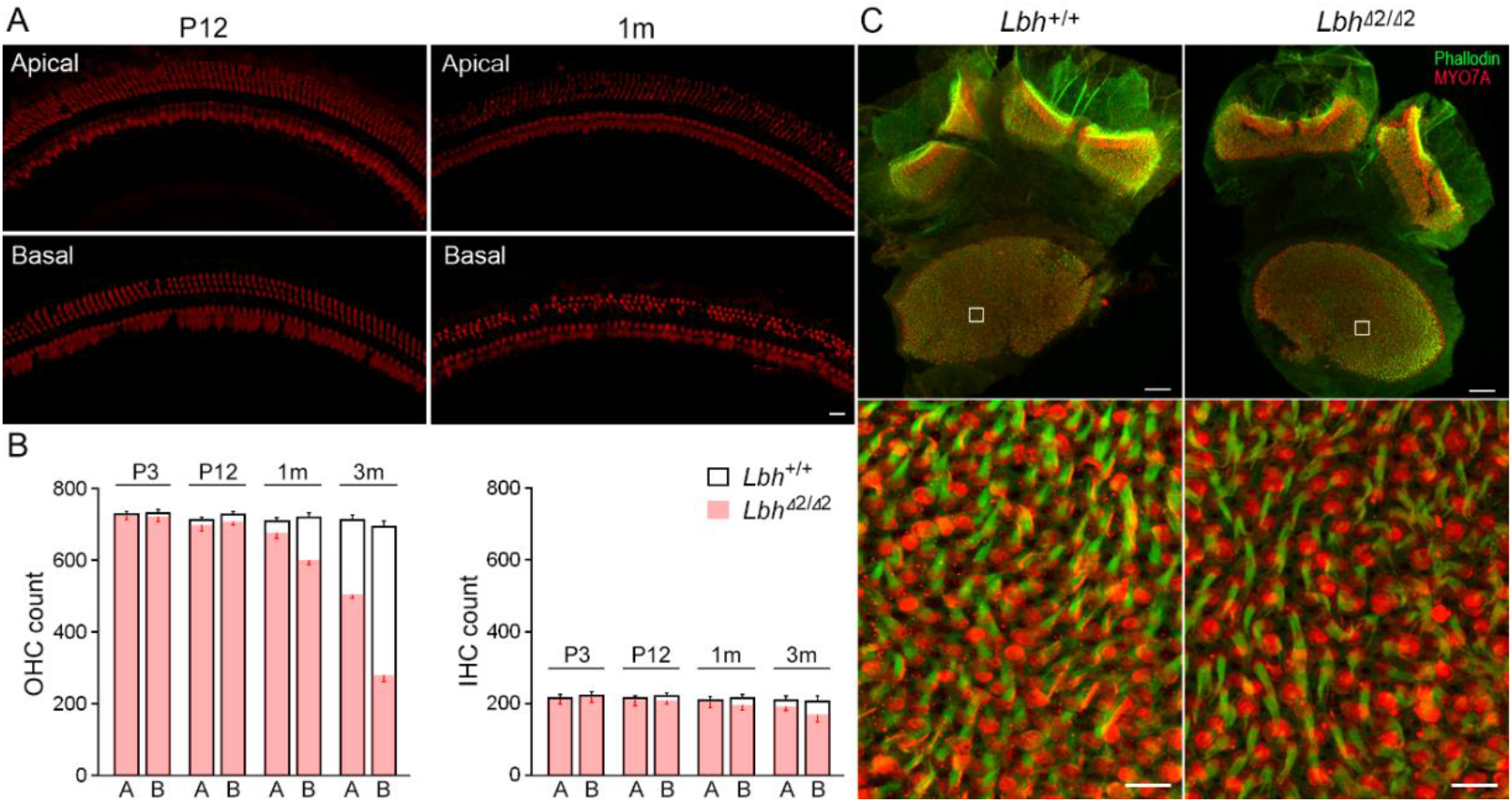
Hair cell status in the cochlear and vestibular sensory epithelia. **A**: Confocal micrographs of IHCs and OHCs labelled by anti-MYO7A-antibody. Images were obtained from an apical and a basal region in the cochleae of *Lbh*^Δ2/Δ2^ null mice at P12 and 1 month. Bar: 10 μm for all panels in A. **B**: IHC and OHC count (mean ± SD) from apical (A) and basal (B) regions of four *Lbh*^Δ2/Δ2^ (pink color) and four *Lbh*^+/+^ (black lines) mice at P3, P12, 1 month and 3 months. **C**: Utricle and crista ampulla of *Lbh*^+/+^ and *Lbh*^Δ2/Δ2^ mice at 3 months (top panel). Bar: 50 μm. Higher magnification images of areas within white frames are presented in the bottom panels. Bar: 10 μm.

Scanning electron microscopy (SEM) was used to examine stereocilia bundle morphology in *Lbh*^Δ2/Δ2^ mice to determine whether LBH is necessary for morphogenesis and maintenance of stereocilia and for differentiation of IHCs and OHCs. Fig. 4A shows an electron micrograph of stereocilia bundles in a P5 *Lbh*^Δ2/Δ2^ mouse cochlea. The characteristic one row of IHC and three rows of OHC stereocilia bundles were well organized and properly oriented. At higher magnification (Figs. 4B,C), the stereocilia were arranged in a normal staircase fashion, with OHCs (Fig. 4B) and IHCs (Fig. 4C) having distinct morphologies. Thus, no signs of abnormality or degeneration of the stereocilia bundles were visible at P5. In one-month-old *Lbh*^Δ2/Δ2^ cochleae, stereocilia bundles in the apical turn appeared largely normal, although sporadic OHC stereocilia bundle loss was observed (asterisk in Fig. 4D). However, degeneration and loss of OHC stereocilia bundles in the basal turn were more pronounced (Fig. 4E). Some of the remaining bundles showed signs of degeneration such as absorption (marked by arrows in Fig. 4F), corruption and recession of the stereocilia on the edge of the bundle (Fig. 4G). While the majority of IHC stereocilia bundles in the basal turn were present (Fig. 4E), some signs of IHC degeneration (such as fusion of the stereocilia in Fig. 4H) were also observed. The fact that the stereocilia bundles of IHCs and OHCs looked normal in P5 *Lbh*^Δ2/Δ2^ mice suggests that LBH is not essential for morphogenesis of the stereocilia bundles. However, degeneration and loss of stereocilia bundles in adult *Lbh*^Δ2/Δ2^ HCs suggest that the maintenance of the bundles and survival of HCs, especially OHCs, depend on LBH.

**Figure 4:**
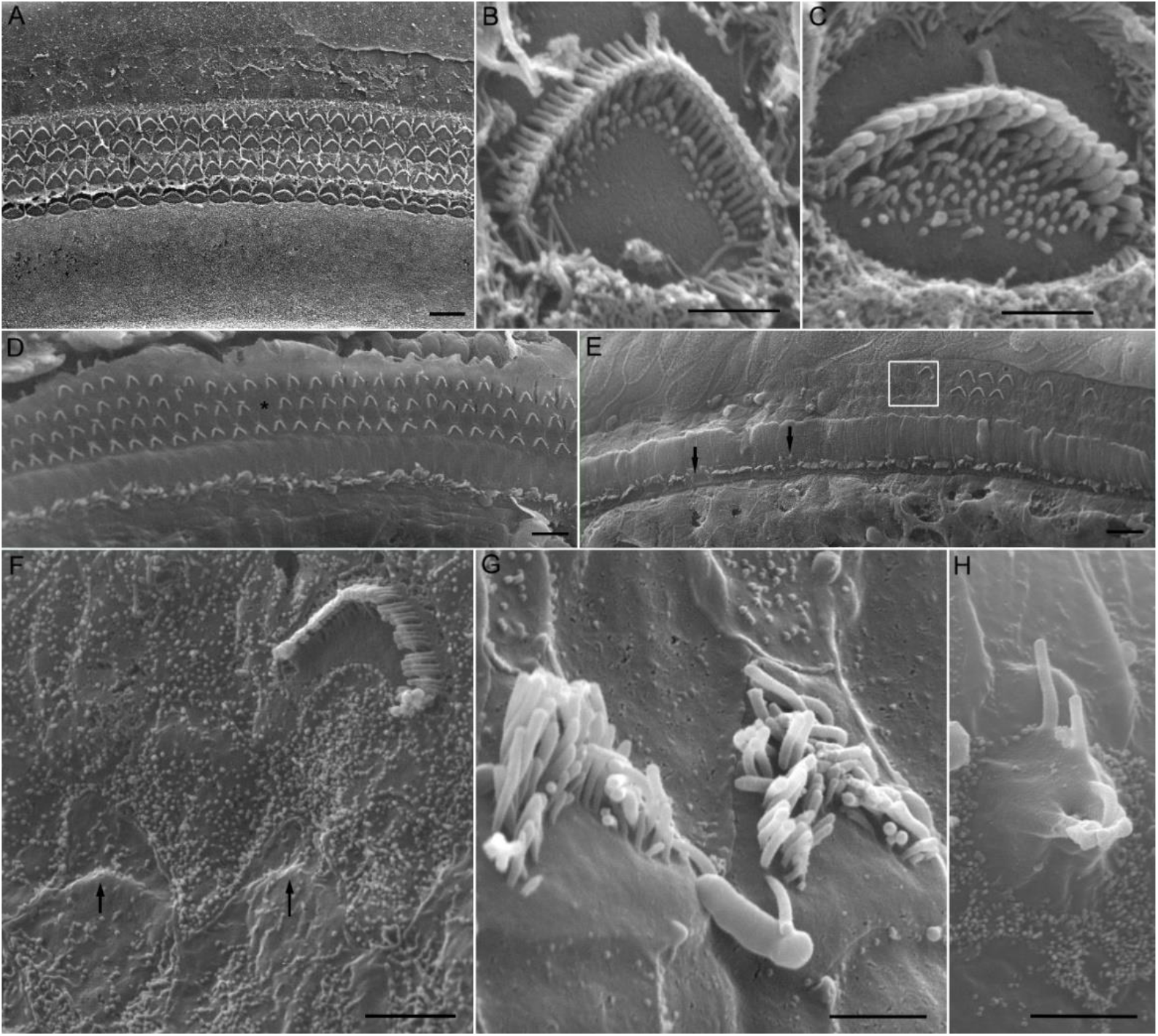
SEM micrographs of stereocilia bundles of cochlear hair cells in *Lbh*^Δ2/Δ2^ null mice. **A**: Micrograph of stereocilia bundles from the low-apical region of a cochlea at P5. Bar: 10 μm. **B** and **C**: Higher magnification images of the stereocilia bundle of an OHC (B) and an IHC (C) from the basal turn of the same cochlea shown in panel A. Bar: 1 μm for B and C. **D** and **E**: Micrographs of stereocilia bundles from an apical turn region (D) and basal turn region (E) from an 1 month-old *Lbh*^Δ2/Δ2^ mouse. Asterisk marks a missing OHC while black arrows mark fusion of stereocilia. Bar: 10 μm. A magnified image of the area within the white frame is highlighted in panel F. **F**, **G** and **H**: Representative images of degenerating stereocilia bundles of OHCs (F,G), and an IHC (H) from mid-basal turn region of an 1-month-old *Lbh*^Δ2/Δ2^ mouse. Black arrows in panel F indicate near complete absorption of stereocilia bundles. Bars: 2 μm (F), 1.5 μm (G), and 2.5 μm (H).

### 4. Mechanotransduction (MET) and electromotility of OHCs in *Lbh*^Δ2/Δ2^ mice

We questioned if LBH plays a role in development of MET apparatus as LBH expression appeared to be limited to HCs. Voltage-clamp technique was used to measure MET current of OHC stereocilia bundles in response to bundle deflection in *Lbh*^Δ2/Δ2^ mice. A coil preparation from the mid-cochlear region was used for recording (Jia and He, 2005). The bundle was deflected with the fluid jet technique (Kros et al., 1992, Jia et al., 2009) and the deflection-evoked MET current was recorded (Fig. 5A). Two examples of the maximal MET current from OHCs of *Lbh*^Δ2/Δ2^ and *Lbh*^+/+^ mice at P12 are presented in Fig. 5A. We compared maximal MET currents from 9 and 8 OHCs from four *Lbh*^+/+^ and four *Lbh*^Δ2/Δ2^ mice, respectively. The magnitude of the current was 614 ± 90 pA (mean ± SD) for *Lbh*^+/+^ and 449 ± 57 pA for *Lbh*^Δ2/Δ2^ OHCs. Despite significant reduction (p=0.00048), the presence of MET current suggests that the mechanotransduction apparatus is functional in *Lbh*^Δ2/Δ2^ OHCs.

**Figure 5:**
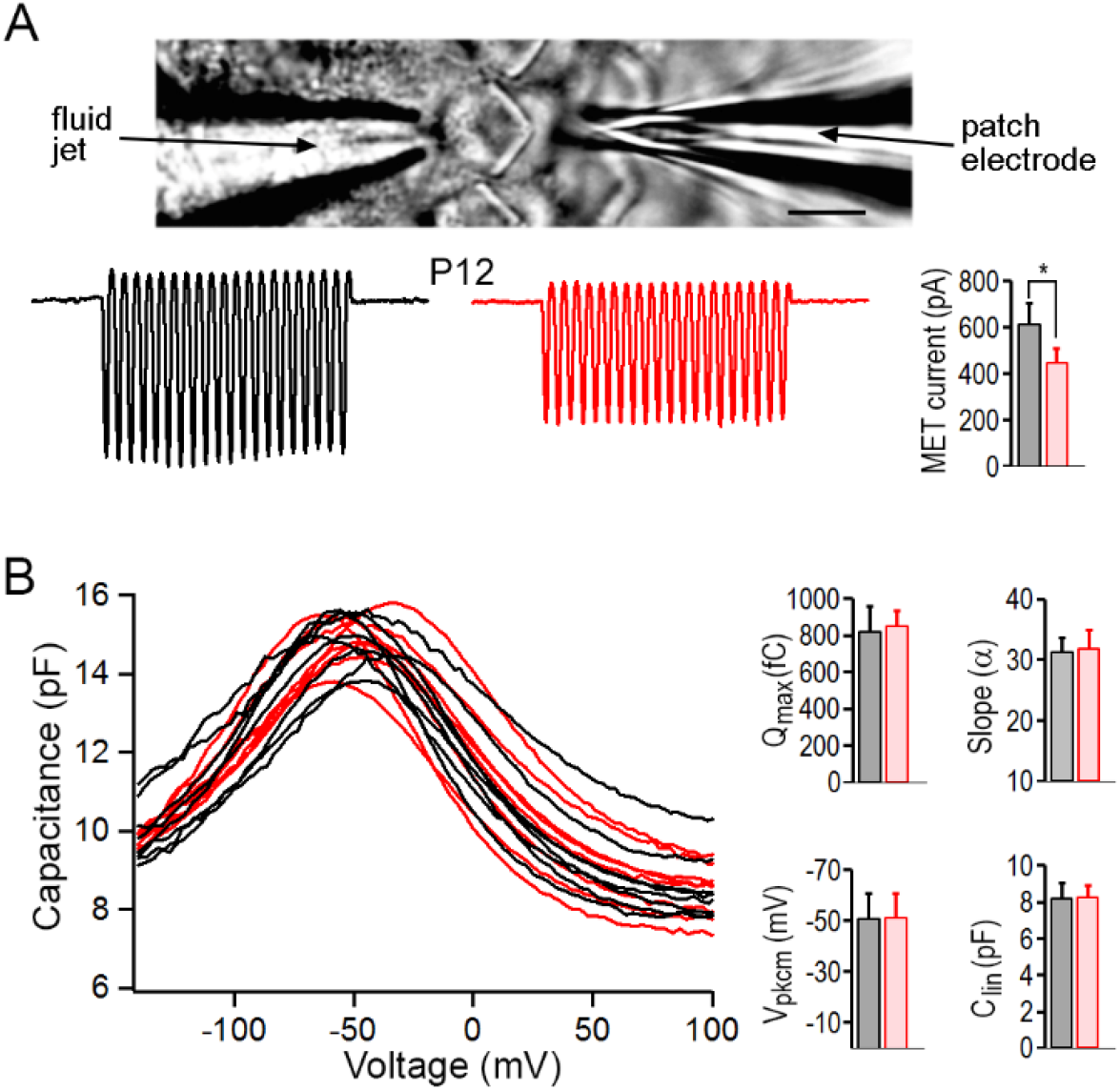
OHC function examined using whole-cell voltage-clamp technique. **A**: Recording of MET current *in vitro* and representative MET current recorded from OHCs in lower apical turn of *Lbh*^Δ2/Δ2^ (red) and *Lbh*^+/+^ (black) mice at P12. Means and SDs are plotted in the right panel. Asterisk marks statistical significance (p<0.05). Bar: 5 μm. **B**: NLC measured from 9 and 8 OHCs in the lower apical turn of *Lbh*^Δ2/Δ2^ (red) and *Lbh*^+/+^ (black) mice, respectively, at P12. Curve fitting using a two-states Boltzmann function yielded four parameters: Q_max_, slope (α), V_pkcm_, and C_lin_. The means and SDs from the two types of OHCs are plotted in the right panels. Student’s *t*-tests yielded p = 0.29, 0.33, 0.47 and 0.42, respectively, for the four parameters.

Prestin-based somatic motility is a unique property of OHCs (Zheng et al., 2000). As LBH is predominantly expressed in OHCs, we asked if LBH regulates prestin expression. OHC electromotility occurs after birth (He et al., 1994; He, 1997); thus, we measured nonlinear capacitance (NLC), an electric signature of electromotility (Ashmore, 1989; Santos-Sacchi, 1991; He et al., 2010), from *Lbh*^Δ2/Δ2^ OHCs at P12 when OHC degeneration was observed to be mild. Fig. 5B shows NLC measured from 9 OHCs from the mid-cochlear region in *Lbh*^Δ2/Δ2^ and *Lbh*^+/+^ mice. A two-state Boltzmann function relating nonlinear charge movement to voltage (Ashmore, 1989; Santos-Sacchi, 1991) was used to compute four parameters, the maximum charge transferred through the membrane’s electric field (Q_max_), the slope factor of the voltage dependence (α), the voltage at peak capacitance (V_pkcm_), and the linear membrane capacitance (C_lin_). No statistically significant differences in any of these parameters were found between *Lbh*^Δ2/Δ2^ and *Lbh*^+/+^ OHCs (Fig. 5B). Thus, OHC motility is not affected by loss of LBH.

### 5. Changes in OHC gene expression after deletion of *Lbh*

To identify molecular mechanism underlying the observed hearing and HC loss in *Lbh*-deficient mice, we performed OHC-specific RNA-seq transcriptome analyses. OHCs were isolated from P12 mice *Lbh*^Δ2/Δ2^ and *Lbh*^+/+^ mice (Fig. 6A), as at this stage HC degeneration in *Lbh* null mice had just begun (Fig. 3A,B). The raw data of transcriptomes of P12 OHCs from *Lbh*^Δ2/Δ2^ and *Lbh*^+/+^ mice are available from the National Center for Biotechnology Information BioProject’s metadata (<u>PRJNA552016</u>). For similarity comparison, Fig. 6B shows a Euclidean distance heatmap of 10,000 genes with a cutoff Z-score calculated as the absolute values from the mean. Comparison of the gene expression profiles between *Lbh*^Δ2/Δ2^ and *Lbh*^+/+^ OHCs identified 2,779 differentially upregulated and 2,065 differentially downregulated genes (defined as those whose expression levels were ≥ 1.0 log2 fold change in expression between the two cell types with statistical significance (FDR *p value* ≤ 0.01)) in *Lbh*^Δ2/Δ2^ OHCs (Fig. 6C). Among those genes, biological processes related to gene expression, protein metabolic process, and organelle organization were significantly enriched in *Lbh*^Δ2/Δ2^ OHCs, as assessed by ShinyGO analysis (Figs. 6D,E). In contrast, *Lbh*^+/+^ OHCs showed greater enrichment in genes associated with cytoskeletal and actin filament organization, membrane bound cell projection organization, anatomical structure morphogenesis, RNA splicing, and axon ensheathment (Fig. 6E).

**Figure 6:**
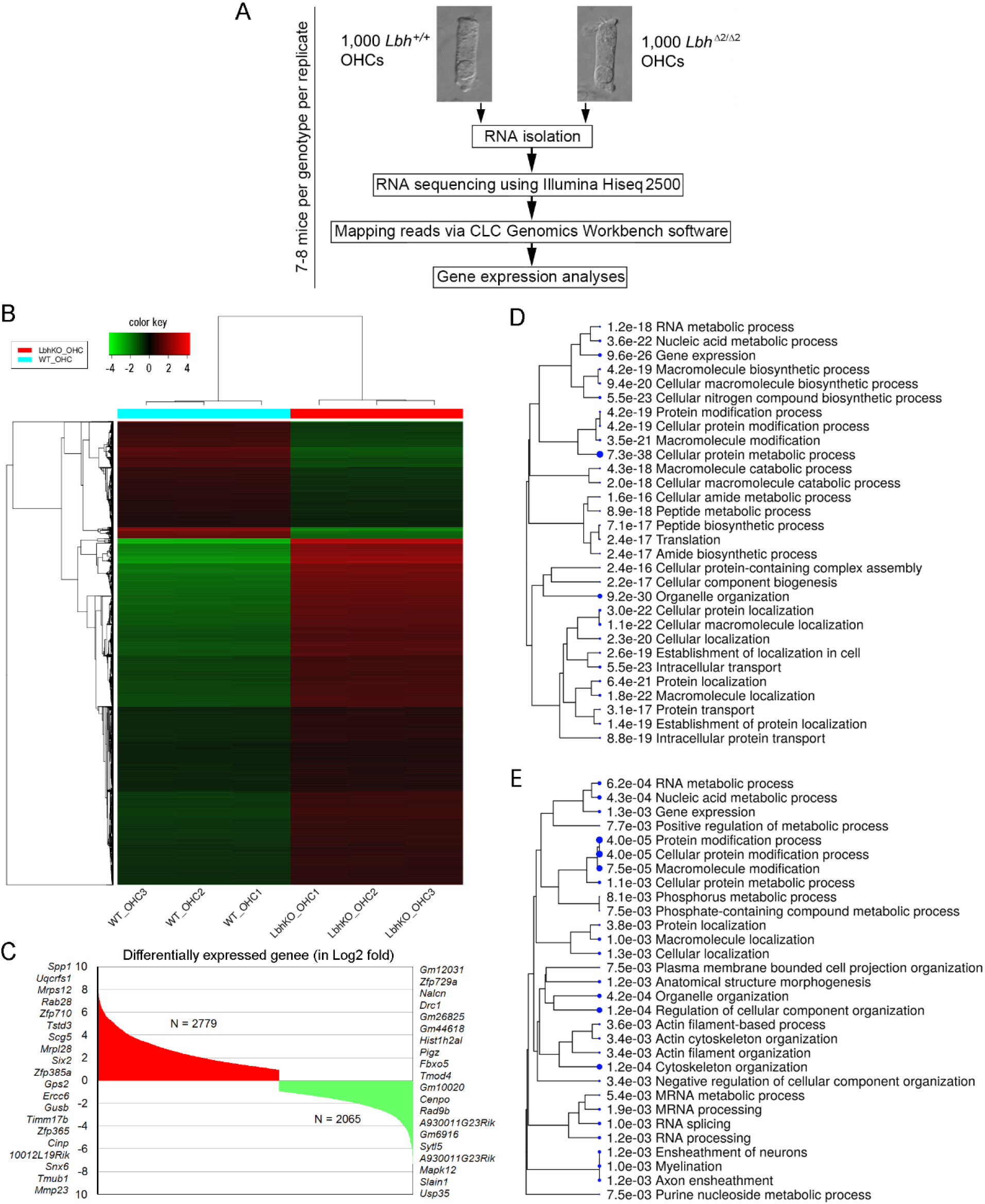
RNA-seq transcriptome analysis of *Lbh*^Δ2/Δ2^ and *Lbh*^+/+^ OHCs. **A**: Workflow of the experimental design for RNA-seq analysis of OHCs isolated from *Lbh*^Δ2/Δ2^ and *Lbh*^+/+^ mice. **B**: Euclidean distance heatmap of 10,000 genes (Z-score cutoff = 4), depicting average linkage between genes expressed in *Lbh*^Δ2/Δ2^ and *Lbh*^+/+^ OHCs. **C**: Upregulated and downregulated genes in *Lbh*^Δ2/Δ2^ compared to *Lbh*^+/+^ OHCs. The top 20 genes up- or down-regulated are shown on either side of the plot. **D**: ShinyGO biological processes enriched in upregulated genes in *Lbh*^Δ2/Δ2^ compared to *Lbh*^+/+^ OHCs. **E**: ShinyGO biological processes enriched in downregulated genes in *Lbh*^Δ2/Δ2^ compared to *Lbh*^+/+^ OHCs.

Additionally, gene set enrichment analysis (GSEA) was performed using the Broad Institute software. Enriched pathways in *Lbh*^Δ2/Δ2^ compared to *Lbh*^+/+^ OHCs included Wnt and Notch signaling pathways, as well as cell cycle regulation, regulation of nucleic acid-templated transcription, DNA damage/repair and autophagy (Fig. 7). As Wnt and Notch play important roles in HC differentiation and regeneration, and LBH is a Wnt target gene known to regulate cell differentiation states in other cell types)(Briegel et al., 2005; Conen et al., 2009; Rieger et al., 2010; Lindley et al, 2015; Li et al., 2015), the expression of genes related to Wnt and Notch signaling was examined in more detail. While *Notch1* (although not among the top 20), *Wnt4*, *Fzd4*, *Ctnnb1* (β-catenin) and *Fuz* were all significantly upregulated, key target genes of Wnt (e.g. *Axin2, Lgr5, Lrp6*) and Notch (i.e. *Hey1, Hey2*), that mirror signaling activity, were downregulated in *Lbh*^Δ2/Δ2^ OHCs (Fig. 7A,B). For cell cycle control, 192 genes were upregulated while 107 were downregulated (Fig. 7C). Analysis of transcription factors shows 430 and 281 transcription factors up- or down-regulated in the *Lbh*^Δ2/Δ2^ OHCs, respectively (Fig. 7D). As LBH is implicated in DNA damage/repair in some cells (Deng et al., 2010; Matusda et al., 2017), the enrichment in these genes was also analyzed (Figs. 7E,F). 90 and 55 genes associated with DNA damage/repair are up- and down-regulated in *Lbh*^Δ2/Δ2^ OHCs, respectively. Interestingly, autophagy-related genes were also found to be enriched, whereby 81 genes were upregulated and 46 downregulated in *Lbh*^Δ2/Δ2^ OHCs.

**Figure 7:**
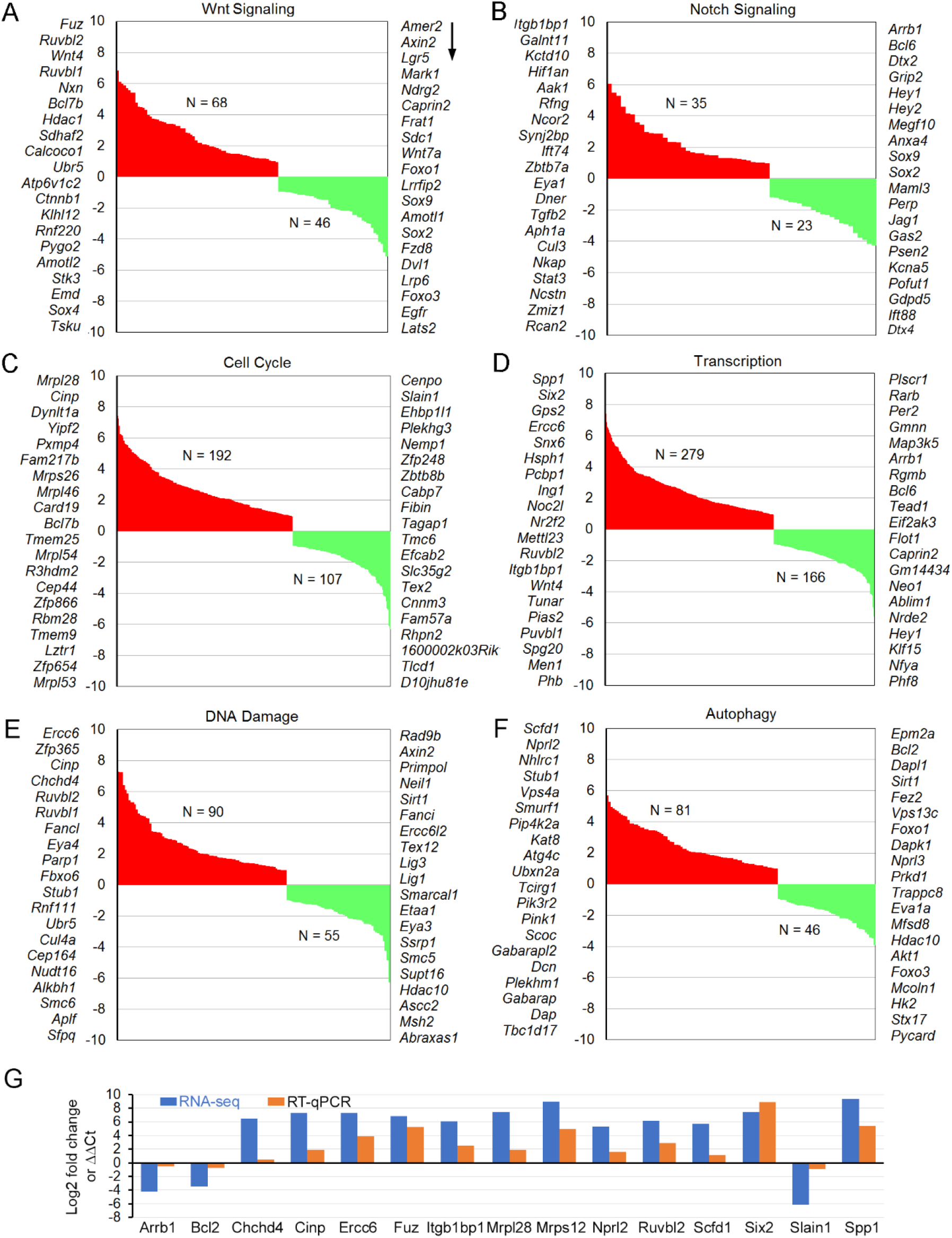
Gene set enrichment analysis (GSEA) of *Lbh*^Δ2/Δ2^ and *Lbh*^+/+^ OHCs transcriptomes. Enriched pathways (FDR < 0.25) *Lbh*^Δ2/Δ2^ null OHCs include regulation of Wnt signaling (A), Notch signaling (B), cell cycle (C), nucleic acid-templated transcription (D), DNA damage/repair (E), and autophagy (F). The total numbers of up-(red) and down-(green) regulated genes within each pathway are indicated, and the top 20 genes in each category are listed on either side of the graph, with greatest to least fold change in downward direction (arrow). G: Validation of differentially expressed cell survival genes using RT-qPCR. Log2 fold changes (*Lbh*^Δ2/Δ2^ *vs. Lbh*^+/+^) from RNA-seq and ΔΔCt values (normalized to *Gapdh*) from RT-qPCR for each gene are shown.

RT-qPCR was used to validate selected differentially expressed genes identified by the RNA-seq analysis, using RNA from P12 *Lbh*^Δ2/Δ2^ and *Lbh*^+/+^ OHCs. Seventeen genes involved in key biological processes related to HC maintenance/degeneration were chosen for comparison. As shown in Fig. 7G, the trend of differential expression of these genes is highly consistent between the two analyses, confirming LBH-dependent gene expression changes in the global RNA-seq analysis.

## Discussion

LBH, a transcriptional regulator highly conserved in evolution from zebrafish to human, is implicated in heart (Briegel and Joyner, 2001; Briegel et al., 2005; Ai et al. 2008), bone (Conen et al. 2009), and mammary gland (Lindley et al., 2015) development. A zebrafish LBH homologue, *lbh-like*, is necessary for photoreceptor differentiation (Li et al., 2015). Here, we identified an unanticipated novel role of LBH in the maintenance of the adult auditory sensory epithelium.

Unlike in heart, bone, mammary gland, and eye development, where LBH or *lbh-like* proteins control progenitor/stem cell fate, self-renewal, and/or differentiation, LBH does not appear to be critical for cochlear HC differentiation, specification, and stereocilia morphogenesis. Morphologically distinct IHCs and OHCs were present at birth, with no HC loss in the cochleae of *Lbh*-null mice at early postnatal stages. Stereocilia bundles of *Lbh*-null HCs also appeared normal and were functional, as mechanical stimulus was able to evoke MET currents. Furthermore, LBH is not necessary for expression of prestin, a specialization of OHCs, despite the fact that *Lbh* is preferentially expressed in adult OHCs. We, therefore, conclude that LBH is not necessary for stereocilia morphogenesis and HC differentiation, specification and development.

Surprisingly, however, we found that LBH is critical for stereocilia bundle maintenance and survival of HCs in adult mice. When *Lbh* was deleted, stereocilia and HCs began to degenerate as early as P12. The degeneration was progressive from OHCs to IHCs and from base to apex of the cochlea, similar to the pattern seen during age-related hearing loss. Our findings, to the best of our knowledge, are the first demonstration that loss of LBH causes degeneration of cochlear HCs, leading to progressive hearing loss. Furthermore, these results provide evidence that LBH is required for adult tissue maintenance. While LBH has been previously implicated in tissue maintenance and regeneration of the postnatal mammary gland by promoting the self-renewal and maintenance of the basal mammary epithelial stem cell pool (Lindley et al., 2015); HCs are terminally differentiated, postmitotic cells that have lost the ability to proliferate and regenerate. In this regard, it is worth noting that loss of LBH is also associated with Alzheimer’s, a neurodegenerative disease affecting postmitotic neurons (Yamaguchi-Kabata et al., 2018). Thus, LBH appears to be required for tissue maintenance in both regenerative and non-regenerative adult tissues.

OHC-specific RNA-seq and bioinformatic analyses examined the potential molecular mechanisms underlying HC degeneration after *Lbh* deletion. Our analyses showed that a greater number of genes were upregulated in *Lbh-*null OHCs compared to wildtype littermate OHCs. Importantly, genes/pathways associated with transcriptional regulation, cell cycle, DNA repair/maintenance, autophagy, as well as Wnt and Notch signaling were significantly enriched in *Lbh*-null OHCs. Notch and Wnt signaling are known to be critical for differentiation and specification of HCs and supporting cells during inner ear morphogenesis and development (Raft and Groves, 2015). Notch and Wnt signaling are also important for transdifferentiation of supporting cells to HCs during regeneration (Waqas et al., 2016; Jansson et al., 2015; Zak et al., 2015). Blocking Notch signaling leads to generation of supernumerary HCs *in vivo* and *in vitro* (Lanford et al., 1999; Li et al., 2018). Notch and Wnt signaling are normally downregulated (Kiernan, 2013), whereas *Lbh* is upregulated during HC maturation (Scheffer et al., 2015). Interestingly, our RNA-seq and pathway analyses showed that, although many WNT and Notch pathway genes were upregulated in *Lbh-*null OHCs, key target genes of WNT (*Axin2, Lgr5, Lrp6*) and Notch (*Hey1, Hey2, Jag1*), mirroring endogenous signaling activity, were downregulated. This suggests that normal adult OHCs retain low levels of Notch and WNT signaling for their maintenance. It also suggests that LBH is required for maintaining low level Notch and WNT activity in OHCs, and that dysregulated Notch/Wnt activity following LBH ablation, as measured by altered WNT/Notch target gene expression in *Lbh* null OHCs, may lead to OHC degeneration. Thus, LBH may promote maintenance of HCs and stereocilia bundles by regulating Notch and Wnt signaling activity. Alternatively, OHC degeneration caused by LBH loss may be due to increased genotoxic and cell stress, as cell cycle, DNA repair/maintenance, and autophagy genes were also deregulated. Indeed, a recent study showed that LBH is involved in cell cycle regulation, and LBH-deficiency induced S-phase arrest and increased DNA damage in articular cartilage (Matusda et al., 2017). Cell-based transcriptional reporter assays further indicate LBH may repress the transcriptional activation of p53 (Deng et al., 2010), a key regulator of DNA damage control and apoptosis.

Our finding that Wnt pathway genes were dysregulated in *Lbh-*null OHCs was unexpected. LBH is a direct WNT/ß-catenin target gene induced by canonical Wnt signaling in epithelial development and cancer (Rieger et a., 2010). It is required downstream of WNT to promote mammary epithelial cell proliferation, while blocking differentiation (Rieger et a., 2010; Ashad-Bishop et al., 2019). Interestingly, LBH knockout in a WNT-driven breast cancer mouse model, MMTV-Wnt1, reduced cell proliferation and hyperplasia, but also increased cell death (Ashad-Bishop et al., 2019). This supports our notion that LBH is required for epithelial cell maintenance. Conversely, knockdown of zebrafish *lbh-like* increased cell proliferation and Notch target gene (*Hes5*) expression (Li et al., 2015); while we find essential Notch target effectors (*Hey1, Hey2*) to be downregulated in *Lbh-*null OHCs. Regardless of whether LBH acts upstream or downstream of Wnt and Notch signaling, all these studies suggest that dysregulation of Notch and Wnt signaling, and alterations in LBH levels can perturb the balance between proliferation, differentiation, and maintenance, with different outcomes in different epithelial tissues (Rieger et al., 2010; Li et al., 2015; Ashad-Bishop et al., 2019).

LBH function as a transcription cofactor has been shown by multiple studies ((Briegel and Joyner, 2001; Briegel et al., 2005; Ai et al 2008; Deng et al 2010). Interestingly, while LBH is predominantly localized to the nucleus in most cells (Briegel and Joyner, 2001; Lindley et al., 2015; Liu et al., 2015; Ashad-Bishop et al., 2019), LBH expression was predominantly cytoplasmic in HCs, although weak nuclear LBH positivity was also observed. In fibroblast-like COS-7 cells, co-localization analysis shows that LBH proteins are localized to both the nucleus and the cytoplasm (Ai et al., 2008). In postmitotic neurons, LBH is also found to be more cytoplasmic than nuclear (unpublished observation). Some transcriptional co-factors, such as TAZ/YAP, are detected in the cytoplasm and can translocate into the nucleus upon mechano-stimulation (Low et al., 2014). The STAT (signal transducer and activator of transcription) transcription factors are constantly shuttling between nucleus and cytoplasm irrespective of cytokine stimulation (Meyer and Vinkemeier, 2004). It is therefore plausible that cytoplasmic LBH in OHCs may translocate to the nucleus upon sensory input. It is also possible that in the cytoplasm LBH may interact with different proteins and have a different function than in the nucleus.

Collectively, this is the first study showing that transcription co-factor LBH can influence stereocilia bundle maintenance and survival of cochlear HCs, especially OHCs. Although the underlying mechanisms remain to be further investigated, based on our analyses, we entertain the possibility that dysregulation of Notch and Wnt signaling caused by LBH loss is detrimental to maintenance of stereocilia bundles and survival of adult cochlear HCs. Importantly, our work points to LBH as a novel causative factor and putative molecular target in progressive hearing loss. It also identifies LBH as paramount for adult tissue maintenance, which could be exploited therapeutically to slow aging of HCs.

## Materials and Methods

### 1. *Lbh* knockout mice

*Lbh*-mutant mice aged between P0 and 3 months were used for experiments. Mice with a conditional null allele of *Lbh* were generated by flanking exon 2 with *loxP* sites (*Lbh*^*flox*^) (17) (Lindley and Briegel, 2013). *Lbh*^*loxP*^ mice were then crossed with a *Rosa26-Cre* line, resulting in ubiquitous deletion of exon 2 and abolishment of LBH protein expression, which was confirmed by western plot and negative anti-LBH antibody staining (Lindley and Briegel, 2013). Ubiquitous *Lbh*-null mice are viable and fertile. Care and use of the animals in this study were approved by the Institutional Animal Care and Use Committees of Creighton University and the University of Miami.

### 2. ABR and DPOAE measurements

ABRs were recorded in response to tone bursts from 4 to 50 kHz using standard procedures described previously (Zhang et al., 2013). Response signals were amplified (100,000x), filtered, averaged and acquired by TDT RZ6 (Tucker-Davis Technologies, Alachua, FL). Threshold is defined visually as the lowest sound pressure level (in decibel) at which any wave (wave I to wave IV) is detected and reproducible above the noise level.

The DPOAE at the frequency of 2f_1_ -f_2_ was recorded in response to f_1_ and f_2_, with f_2_/f_1_ = 1.2 and the f_2_ level 10 dB lower than the f_1_ level. The sound pressure obtained from the microphone in the ear-canal was amplified and Fast-Fourier transforms were computed from averaged waveforms of ear-canal sound pressure. The DPOAE threshold is defined as the f_1_ sound pressure level (measured in decibels) required to produce a response above the noise level at the frequency of 2f_1_-f_2_.

### 3. Recording of CM and EP

Procedures for recording CM and EP were described before (Zhang et al., 2014; Liu et al., 2016). A silver electrode was placed on the ridge near the round window for recording CM. An 8 kHz tone burst was delivered through a calibrated TDT MF1 multi-field magnetic speaker. The biological signals were amplified using an Axopatch 200B amplifier (Molecular Devices, Sunnyvale, CA) and acquired by software pClamp 9.2 (Molecular Devices) running on an IBM-compatible computer. The sampling frequency was 50 kHz.

For recording the EP, a basal turn location was chosen. A hole was made using a fine drill. A glass capillary pipette electrode (5 MΩ) was mounted on a hydraulic micromanipulator and advanced until a stable positive potential was observed. The signals were filtered and amplified under current-clamp mode using an Axopatch 200B amplifier and acquired by software pClamp 9.2. The sampling frequency was 10 kHz.

### 4. Immunocytochemistry and HC count

The cochlea and vestibule from the *Lbh*^Δ2/Δ2^ and *Lbh*^+/+^ were fixed for 24 hours with 4% paraformaldehyde (PFA). The basilar member, including the organ of Corti, the utricle and ampulla were dissected out. Antibodies against MYO7A (Proteus, product # 25-6790) or LBH (Sigma, Lot# HPA034669) and secondary antibody (Life Technologies, Lot# 1579044) were used. Tissues were mounted on glass microscopy slides and imaged using a Leica Confocal Microscope (Leica TCS SP8 MP). HC counts from two areas (approximately 1.4 and 4.5 mm from the hook, each 800 μm in length) were obtained for HC count from confocal images off-line (Zhang et al., 2013).

### 5. SEM

The cochleae from *Lbh*-mutant mice were fixed for 24 hours with 2.5% glutaraldehyde in 0.1 M sodium cacodylate buffer (pH 7.4) containing 2 mM CaCl_2_, washed in buffer. After the cochlear wall was removed, the cochleae were then post-fixed for 1 hour with 1% OsO_4_ in 0.1 M sodium cacodylate buffer and washed. The cochleae were dehydrated via an ethanol series, critical point dried from CO_2_ and sputter-coated with gold. The morphology of the HCs was examined in a FEI Quanta 200 scanning electron microscope and photographed.

### 6. Whole-cell voltage-clamp techniques for recording MET current and NLC

Details for recording MET currents from auditory sensory epithelium are provided elsewhere (Kros et al., 1992; Jia and He, 2005). A segment of auditory sensory epithelium was prepared from the mid-cochlear and bathed in extracellular solution containing (in mM) 120 NaCl, 20 TEA-Cl, 2 CoCl_2_, 2 MgCl_2_, 10 HEPES, and 5 glucose at pH 7.4. The patch electrodes were back-filled with internal solution, which contains (in mM) CsCl 140; CaCl_2_: 0.1; MgCl_2_ 3.5; MgATP: 2.5; EGTA-KOH 5; HEPES-KOH 10. The solution was adjusted to pH 7.4 and osmolarity adjusted to 300 mOsm with glucose. The pipettes had initial bath resistances of ~3-5 MΩ. After the whole-cell configuration was established and series resistance was ~70% compensated, the cell was held under voltage-clamp mode to record MET currents in response to bundle deflection by a fluid jet positioned ~10–15 μm away from the bundle. Sinusoidal bursts (100 Hz) with different magnitudes were used to drive the fluid jet as described previously (Kros et al., 1992; Jia and He, 2005). Holding potential was normally set near −70 mV. The currents (filtered at 2 kHz) were amplified using an Axopatch 200B amplifier and acquired using pClamp 9.2. Data were analyzed using Clampfit in the pClamp software package and Igor Pro (WaveMetrics, Inc).

For recording NLC, the cells were bathed in extracellular solution containing (in mM) 120 NaCl, 20 TEA-Cl, 2 CoCl_2_, 2 MgCl_2_, 10 HEPES, and 5 glucose at pH 7.4. The internal solution contains (in mM): 140 CsCl, 2 MgCl_2_, 10 EGTA, and 10 HEPES at pH 7.4. The two-sine voltage stimulus protocol (10 mV peak at both 390.6 and 781.2 Hz) with subsequent fast Fourier transform-based admittance analysis (jClamp, version 15.1) was used to measure membrane capacitance using jClamp software (Scisoft). Fits to the capacitance data were made in IgorPro (Wavemetrics). The maximum charge transferred through the membrane’s electric field (Q_max_), the slope factor of the voltage dependence (α), the voltage at peak capacitance (V_pkcm_), and the linear membrane capacitance (C_lin_) were calculated.

### 7. Cell isolation, RNA preparation, and RNA-sequencing

*Lbh*^Δ2/Δ2^ and *Lbh*^+/+^ mice at P12 were used for gene expression analysis. Details for cell isolation and collection are provided elsewhere (Liu et al., 2014). Approximately 1,000 OHCs were collected from 7-8 mice for one biological repeat per genotype. Three biological replicates were prepared for each genotype.

Total RNA, including small RNAs (> ~18 nucleotides), were extracted and purified using the Qiagen RNeasy mini plus Kit (Qiagen, Germantown, MD). To eliminate DNA contamination in the collected RNA, on-column DNase digestion was performed. The quality and quantity of RNA were examined using an Agilent 2100 BioAnalyzer (Agilent, Santa Clara, CA).

Genome-wide transcriptome libraries were prepared from three biological replicates separately for *Lbh*^Δ2/Δ2^ and *Lbh*^+/+^ OHCs. The SMART-Seq V4 Ultra Low Input RNA kit (Clontech Laboratories, Inc., Mountain View, CA) and the Nextera Library preparation kit (Illumina, Inc., San Diego, CA) were used. An Agilent 2100 Bioanalyzer and a Quibit fluorometer (Invitrogen, Thermo Fisher Scientific) were used to assess library size and concentration prior to sequencing. Transcriptome libraries were sequenced using the HiSeq 2500 Sequencing System (Illumina). Four samples per lane were sequenced, generating approximately 60 million, 100 bp single-end reads per sample. The files from the multiplexed RNA-seq samples were demulitplexed and fastq files were obtained.

The CLC Genomics Workbench software (CLC bio, Waltham, MA, USA) was used to individually map the reads to the exonic, intronic, and intergenic sections of the mouse genome (mm10, build name GRCm38). Gene expression values were normalized as reads per kilobase of transcript per million mapped reads (RPKM). Log fold changes and FDR p-values were calculated, and the dataset was exported for further analysis. The raw data are available from the National Center for Biotechnology Information BioProject’s metadata (<u>PRJNA552016</u>) and the normalized RPKM values are accessible as an Excel file. Transcriptomes and differentially expressed genes as well as significantly enriched genes associated with Wnt and Notch signaling, transcription, cell cycle, DNA damage/repair, and autophagy in the *Lbh*-null OHCs are also provided as **Supplemental File 1.**

### 8. Real-time quantitative PCR for validation

OHCs were collected as described above from fourteen additional *Lbh*^Δ2/Δ2^ mice (aged P12) and 15 age-matched *Lbh*^+/+^ mice for RT-qPCR. Total RNA was isolated using the Qiagen miRNeasy kit and quantified by nanodrop. cDNA libraries were prepared from isolated RNA with the iSCRIPT master mix (BioRad). Oligonucleotide primers were acquired from Integrated DNA Technologies (Coralville, Iowa). The sequences of oligonucleotide primers are shown as follows:

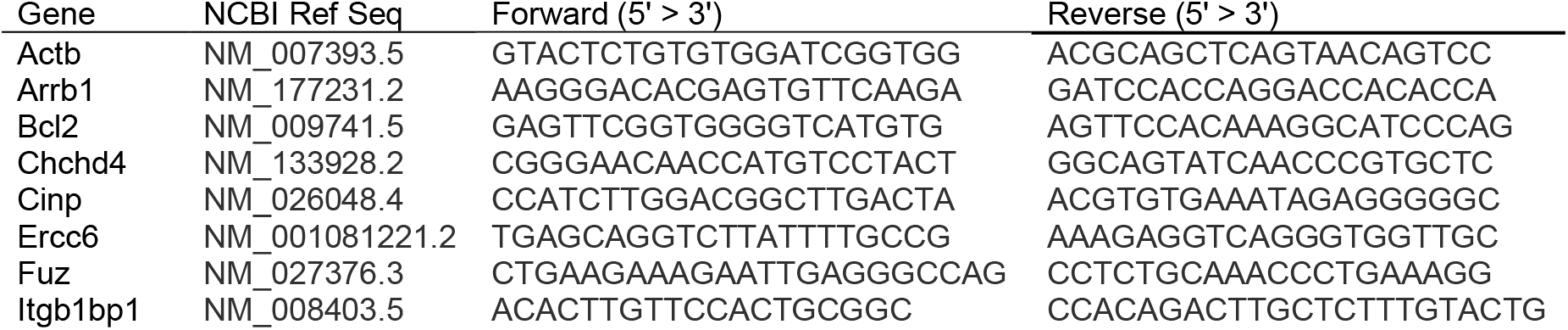
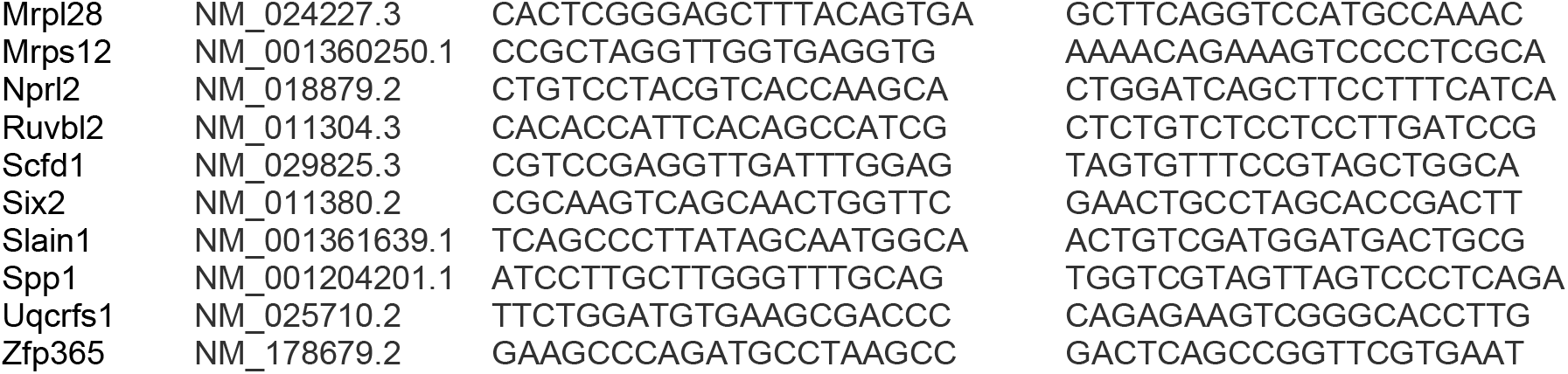

RT-qPCR reactions were prepared as 10 μl reactions including *Lbh*^Δ2/Δ2^ or *Lbh*^+/+^ OHC cDNA, PowerUp SYBR green master mix (Thermo Fisher), gene-specific forward and reverse primers and run in triplicate on a BioRad CFX96 Touch real-time PCR machine. Primer specificity was confirmed by melt curve analysis. Quantified expression (Ct) of each gene (gene of interest or GOI) was normalized to the Ct value of a house-keeping gene (*Actb*) (ΔCt = Ct^(GOI)^ ― Ct^AVG *Actb*^). Then differential expression of the gene between *Lbh*^Δ2/Δ2^ and *Lbh*^+/+^ OHCs was calculated as ΔΔCt (ΔΔCt = ΔCt *Lbh*^Δ2/Δ2^ − ΔCt *Lbh*^+/+^). The relationships between the RNA-seq derived-log2 fold change values and ΔΔCq values from RT-qPCR between *Lbh*^Δ2/Δ2^ and *Lbh*^+/+^ OHCs were compared to confirm trends in expression.

### 9. Bioinformatic Analyses

The expressed genes were examined for enrichment using GSEA v. 3.0 (Broad Institute) (Mootha et al., 2003; Subramanian et al., 2005), iDEP 0.85 and ShinyGO (Ge-lab.org) (Ge et al., 2018). Enriched biological processes and molecular functions, classified according to gene ontology (GO) terms, as well as signaling pathways in the *Lbh*^Δ2/Δ2^ and *Lbh*^+/+^ OHCs were examined (FDR cutoff < 0.05). With the RPKM expression value arbitrarily set at ≥ 0.10 (FDR p-value ≤ 0.05) (Li et al., 2018; Liu et al., 2018), expression values from *Lbh*^Δ2/Δ2^ and *Lbh*^+/+^ OHCs were input into iDEP for analyses and log transformed. All transcripts detected in *Lbh*^Δ2/Δ2^ and *Lbh*^+/+^ OHCs is provided in Supplementary Data 1. For reference and verification, additional resources such as the Ensembl database, AmiGO (http://amigo.geneontology.org/amigo), gEAR (www.umgear.org) and SHIELD (https://shield.hms.harvard.edu/index.html) were also used. No custom code was used in the analysis.

### 10. Statistical analysis

Means and standard deviations (SD) were calculated based on measurements from three different types of mice. For each parameter, student’s t-test was used to determine statistical significance between two different conditions and genotypes. Probability (P) value ≤ 0.01 was regarded as significant. For transcriptome analysis, means and SD were calculated for three biological repeats from *Lbh*^Δ2/Δ2^ and *Lbh*^+/+^ OHCs, with four technical replicates each. ANOVA False Discovery Rate-corrected *p*-values were used to compare average expression (RPKM) values for each transcript and FDR *p* < 0.05 was considered statistically significant.

## Acknowledgments

This work has been supported by the NIH grants R01DC016807 from the NIDCD to DH, R01 GM113256 from the NIGMS to KJB and R01DC005575 and R01DC012115 from the NIDCD to XL. YL is supported by National Science Foundation of China (#81600798 and #81770996). We acknowledge the use of the University of Nebraska DNA Sequencing Core Facility for performing RNA-seq. The University of Nebraska DNA Sequencing Core receives partial support from the NCRR (RR018788).

## Author Contributions

HL performed electrophysiological and morphological experiments, KG performed transcriptome analysis and contributed to manuscript writing, GMH, SM, YL, XL performed some morphological and electrophysiological experiments, KB generated *Lbh*-mutant mice, contributed to experimental design, data analysis, and manuscript writing. DH designed research, performed SEM experiments and wrote the manuscript.

## Other supplementary materials for this manuscript include the following

Datasets (in Excel format): RNA-seq dataset of transcriptomes of *Lbh*-null and wildtype outer hair cells. Differentially expressed genes as well as significantly enriched genes associated with Wnt and Notch signaling, transcription, cell cycle, DNA damage/repair, and autophagy in the *Lbh*-null OHCs are also included.

